# Develope Micro clonal -propagation protocol for *Oxytenanthera abyssinica A.Rich. Munro* to large scale micro-propagation

**DOI:** 10.1101/2020.04.28.063883

**Authors:** Adugnaw Admas, Berhane Kidane, Melaku Admasu, Tesaka Misga

## Abstract

In Ethiopia, *Oxytenanthera abyssinica A.Rich. Munro* has varies economic importance. However, conventional propagation methods of *O. abyssinica* are generally inefficient due to their low multiplication rate, time consuming, labor intensive, and too costly. The objective of this study was to develop a protocol for mass micropropagation of O. abyssinica through seed culture. Murashige and Skoog (MS) medium augmented with 6-Benzylaminopurine (BAP) was used for shoot initiation and multiplication. For in vitro rooting, MS medium supplemented with 3-Indole –butric acid (IBA) was used.

In shoot initiation experiment all viable seeds were proliferated in 5-7 days of culturing. In shoot multiplication at 0.004 g/L BAP was Sucssefuly shoot multiplied, also best root responding were found at 0.005 g/l IBA.

The present optimized protocol enables for any acters who needs large numbers of low land bamboo seedling for industery, small and micro enterprize or for reafforestation programms.

## 1. INTRODUCTION

Bamboo is hardened and fastest-growing perennial grass species [1] and it is a woody culms and gregarious, monocarpic flowering plant [2]. They belong to the subfamily Bambusoideae and family Poaceae (sometimes called Gramineae),in the same family with cereal crops such as rice and wheat and sugar cane [3]. The term bamboo comprises more than 1,500 species that are widely distributed in the tropical, subtropical and temperate regions of all continents except Antarctica and Europe, between 46°N and 47°S. Geographically bamboo distribution can be classified in to three zones: the Asian Pacific zone, the American zone and the Africanzone[4]. The highest diversity and area coverage of bamboo is recorded from the Asian continent, followed by America and Africa [5]. The sizes of bamboos vary from small annuals to giant perennial timber bamboo species [6]. Dwarf bamboos may be as little as 10cm in height, but stands of tall species may attain 15–20m, and the largest known (e.g. Dendro calamus giganteus and Dendrocalamus brandisii) grow up to 40m in height and 30cm in culm (stem) diameter [7; 8; 9].

43 species of bamboo in 11 genera can be found in Africa, covering an estimated area of 3.6 million ha [10]. Out of these African bamboo species, Ethiopia has only two endemic species, namely the highland bamboo (Yushania alpine K. Schumach.) and lowland bamboo (Oxytenanthera abyssinica A.Rich. Munro). These two species are restricted in limited agro ecological regions, i.e. in highland areas of altitude 2400-3500 ma.s.l. and in lowland areas from 500-1800 ma.s.l [11].

For adapation, Ethiopia were imported diffrenet bamboo species and they are under field trial in diffrenet locations those are: *Dendrocalamus asper,Dendrocalamushamiltonii,Dendrocalamusgiganteus,Dendrocalamusmembr anaceusMunro, Bambusa vulgaris Var. green, Bambusa vulgarisVar. Vitata,Guadua amplexifolia* [12].

Bamboo is used for housing, handicrafts, pulp and paper industries, energy source, and food. It has also high value in carbon sequestration [13]. Medical use of O. abyssinica is documented in different countries including Ethiopia [14]. O. abyssinica has also important phytochemicals with a resultant antioxidant property [15]. Furthermore, investigation on bamboo shoots showed that O. abyssinica shoot is rich in nutrients [16].

Conventionally, bamboos are propagated through seeds, clump division, rhizome and culm cuttings [17]. However,gregarious flowering at long intervals followed by the death of clumps, short viability of seeds [18], presence of diseases and some pests [19] are limiting factors to use seeds as valuable source of propagation.

Even vegetative propagation methods have limitation for mass propagation since propagules are difficult to extract, bulky to transport, and planting materials are insufficient in number for large-scale plantation [20]. Considering problems encountered in both sexual and asexual conventional propagation of the O. abyssinica species, inovative method that brings about rapid large scale production of bamboo is highly desirable. In this regard different scholars recommended micropropagation as an excellent means to achieve this aim. The first tissue culture study on bamboo (Dendrocalamus strictus) was conducted by Alexander and Rao [21] who germinated embryos in vitro. Since then different researchers have been publishing scientific articles on successful micropropagation protocol through seed culture in different bamboo species; like, Arya et al. on Dendrocalamus asper [22], Arya et al. on Dendrocalamus hamiltonii [23], and Devi et al. on Dendrocalamus giganteus [24]. Nevertheless, their results show there is an interaction of species with hormonal types and levels included in the culture medium which necessitate the development/optimization of micropropagation protocols for every species Under different conditions. And also, Kahsay et al.; 2017 for O. abyssinica species developed protocol for mass proapagation from seed culture by using 3-BAP,NAA and IBA hormone at different concentation and reproducible protocol that can enable the in vitro rapid multiplication of O. abyssinica from seed culture. The main objective of this paper was, therefore, to develop a protocol for in vitro multiplication of O. abyssinica species from seed culture using 3-BAP and IBA hormone for better improvements of Kahsay *et al*.; 2017. The specfice objectives for this study were to determine, identify an appropriate cytokinin and determine its optimal concentration for shoot proliferation and multiplication; identify an appropriate auxin and determine its optimal concentration for root induction.

## 2. MATERIALS AND METHODS

### 2.1 Source of Experimental Material

The seeds for this study were obtained from Bahradr Enviroment and Forest Research centere, Ethiopia. Healthy seeds were selected carefully and they were stored in plastic bag at +4°C in refrigerator. Seeds were stored more than a year in Bahradr Tissue culture laboratory.

### 2.2 Explants Surface Disinfection

Selected healthy seeds as shown in **figure 2** were sterilized to get ride –off all micro –organisms. Also, the seeds were washed with tape water to remove debris. Then, to get clean seeds it soaked in distilled water for 2 hrs by shaking and washed by double distilled water (DDW) with liquid soap with 2-3 drops of Tween −20 for 20 minutes. Then, treated by antifungal of mancozine 20 g/l for 20 minute and washed the seeds with DDw three times. By follwing those porocedure, seeds were treated by 2% of NaOCl for 20 minute and washed the seeds three times by 2-3 drop of Tween-20 for five minute. After pre-treatment,the seeds were treated with 1% NaOCl and washed three times by DDw. Finally, it treated with 70% ethanol for 30 seconds under laminar air flow cabinet. After sterilization of the MS medium, for shoot initiation three jars for each treatments (0.003, 0.004 and 0.005 g/L BAP) five seeds were placed randomly in completely randomized design (CRD) arrangement.

**Figure 1.**
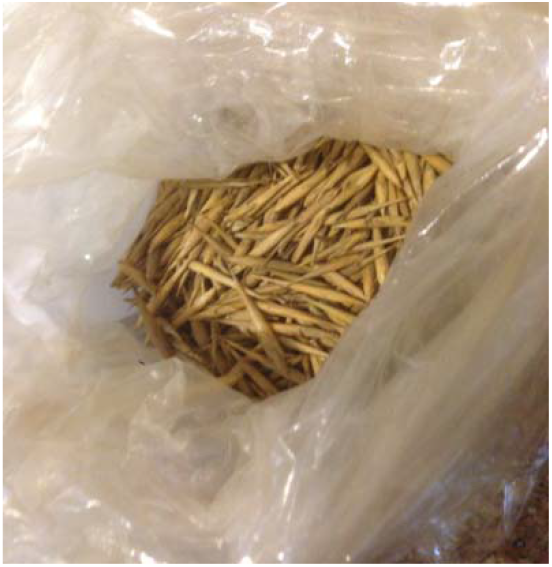
Seeds of *Oxytenanthera abyssinica*

**Figure 2.**
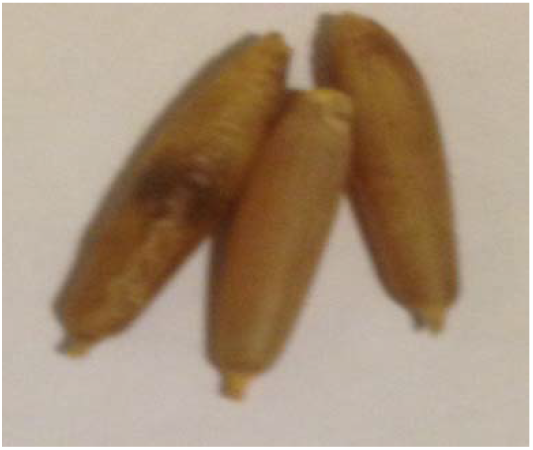
selected healthy seed

### 2.3. Preparation of Stock Solutions for initiation, multiplication and rooting

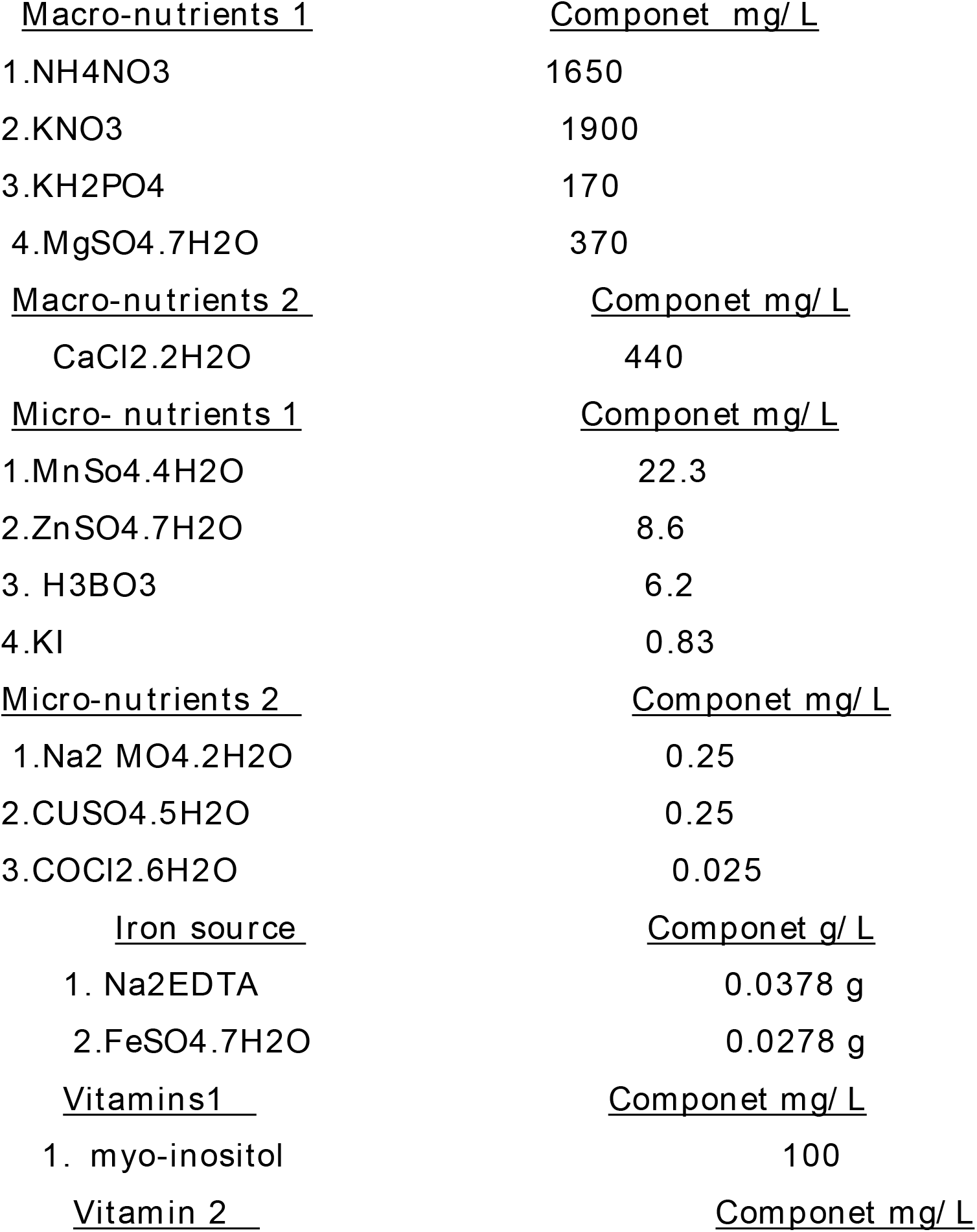
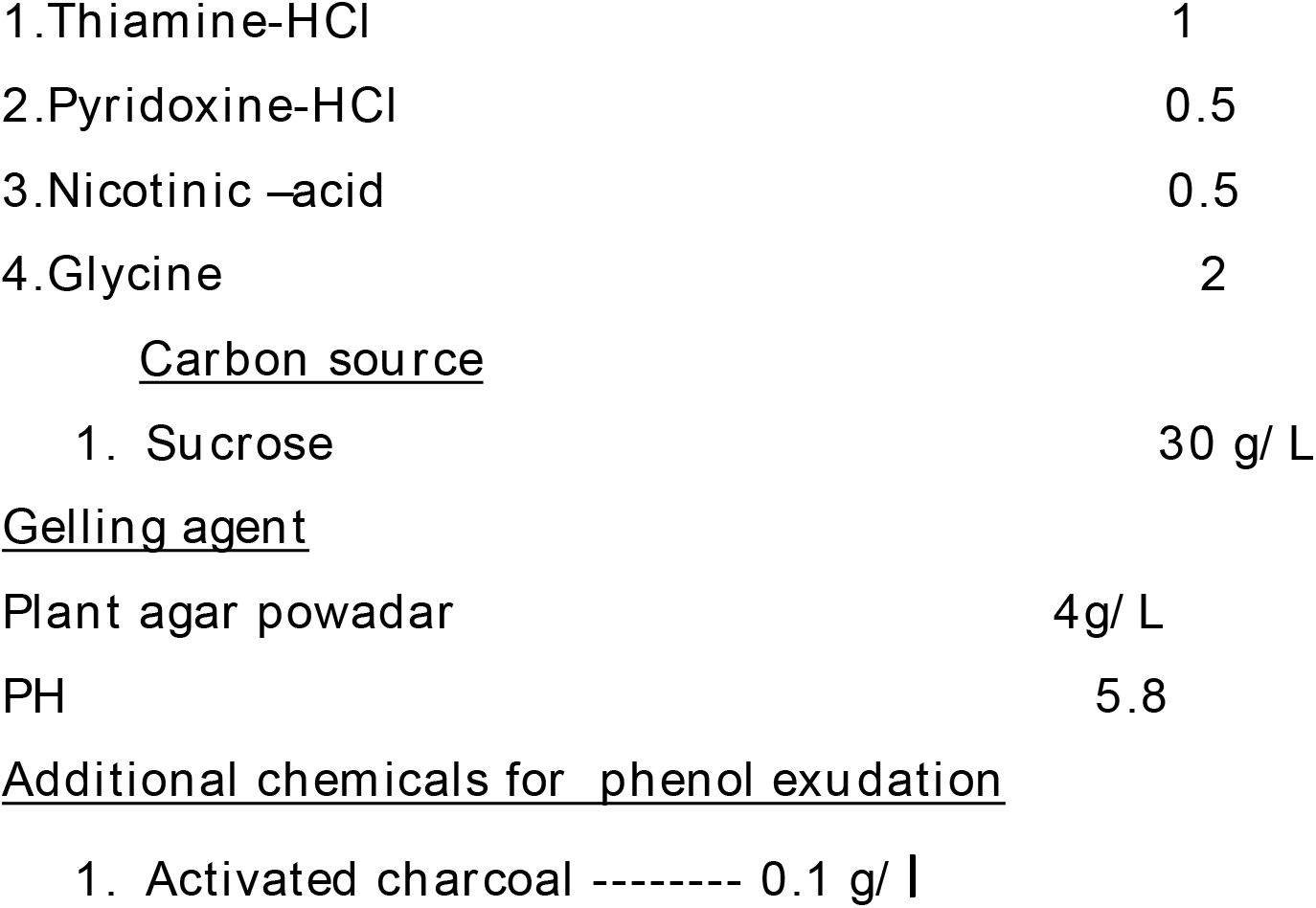

| <u>Macro-nutrients 1</u> | <u>Componet mg/L</u> |
| --- | --- |
| 1.NH<sub>4</sub>NO<sub>3</sub> | 1650 |
| 2.KNO<sub>3</sub> | 1900 |
| 3.KH<sub>2</sub>PO<sub>4</sub> | 170 |
| 4.MgSO<sub>4</sub>.7H<sub>2</sub>O | 370 |
| <u>Macro-nutrients 2</u> | <u>Componet mg/L</u> |
| CaCl<sub>2</sub>.2H<sub>2</sub>O | 440 |
| <u>Micro- nutrients 1</u> | <u>Componet mg/L</u> |
| 1.MnSo<sub>4</sub>.4H<sub>2</sub>O | 22.3 |
| 2.ZnSO<sub>4</sub>.7H<sub>2</sub>O | 8.6 |
| 3. H<sub>3</sub>BO<sub>3</sub> | 6.2 |
| 4.KI | 0.83 |
| <u>Micro-nutrients 2</u> | <u>Componet mg/L</u> |
| 1.Na<sub>2</sub> MO<sub>4</sub>.2H<sub>2</sub>O | 0.25 |
| 2.CUSO<sub>4</sub>.5H<sub>2</sub>O | 0.25 |
| 3.COCl<sub>2</sub>.6H<sub>2</sub>O | 0.025 |
| <u>Iron source</u> | <u>Componet g/L</u> |
| 1. Na<sub>2</sub>EDTA | 0.0378 g |
| 2.FeSO<sub>4</sub>.7H<sub>2</sub>O | 0.0278 g |
| <u>Vitamins1</u> | <u>Componet mg/L</u> |
| 1. myo-inositol | 100 |
| <u>Vitamin 2</u> | <u>Componet mg/L</u> |
| 1.Thiamine-HCl | 1 |
| 2.Pyridoxine-HCl | 0.5 |
| 3.Nicotinic -acid | 0.5 |
| 4.Glycine | 2 |
| Carbon source |  |
| 1. Sucrose | 30 g/L |
| Gelling agent |  |
| Plant agar powder | 4g/L |
| PH | 5.8 |

The above Stock solutions of major and minor salts, vitamins and plant growth regulators (PGRs) of 6-Benzylaminopurine (BAP) acid and Indole-3-butyric (IBA) were used by dissolving in distiled water.

Finally, after adjusting the final volume, the hormone stock solutions were stored in refrigerator at +4°C and other nutrient medium were kept at room temprature and used after 5 days from preparation time for trargeted experment.

### 2.4 Culture Media Preparation

For all experiments the pH of the above mentioned prepared nutrient medium was adjusted to 5.80 before adding 0.4% agar. Full-strength MS medium with 3% sucrose was used for culture initiation and multiplication experiments. About 50 ml of the medium were dispensed to 300 ml jar for initiation experiment and also 50 ml of the medium were dispensed to 300 ml jar for multiplication and rooting experiments. The media were autoclaved at 121°C with 15PSi pressure for 20 minutes and kept under room temperature for four days before used.

### 2.5. Establishment of Culture Shoots

Disinfected seeds were cultured in 9 jars that contined the above mentioned nuterint 50 ml of MS medium with BAP and 3 jars PGRs free medium for shoot initiation study. Then the cultured sample were brought to growth room shelf with a photoperiod of 16/8h light/dark using cool-white fluorescent lamps (photon flux density, 40μmol m-2 s-1 irradiance) at 25 ± 2°C. After seven days, all of the cultured samples were initiated shoot.

### 2.6 Shoot Multiplication

To avoid the carry over effect of shoot initiation media during shoot multiplication, initiated propagules consisting of three shoots each were subcultured on PGRs free MS medium for two weeks. Each propagule was placed vertically and lightly pressed into the culture medium supplemented with of 0.003-0.005 g/litre of BAP with each activated charkol for inhibition of oxidants of the cells mostlly for phenol exudation. MS medium without PGRs was used as control. 12 jars each with three propagules were used and kept under light conditions. Then, after two weeks multipilication of new leaf were best at 0.004 g/litre of BAP as it showed in **figure 5**.

**Figure 3.**
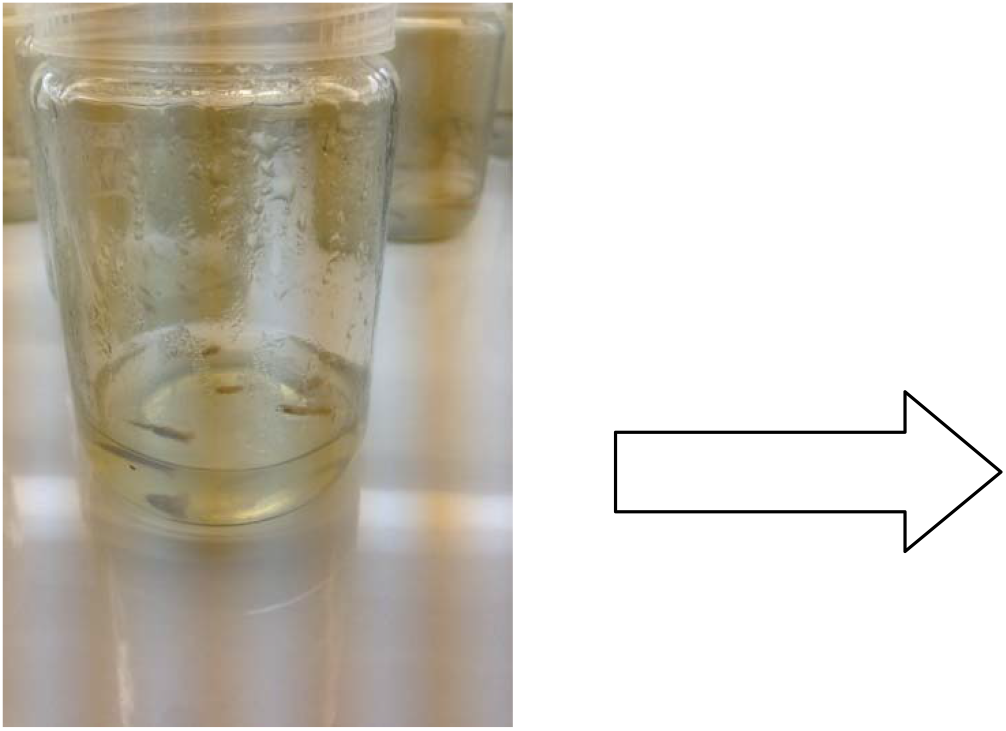
Cultured seed

**Figure 4.**
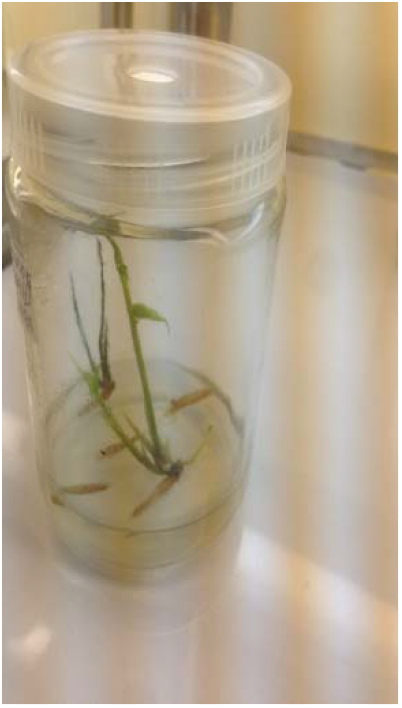
Intiated leaf from cultured seed

**Figure 5.**
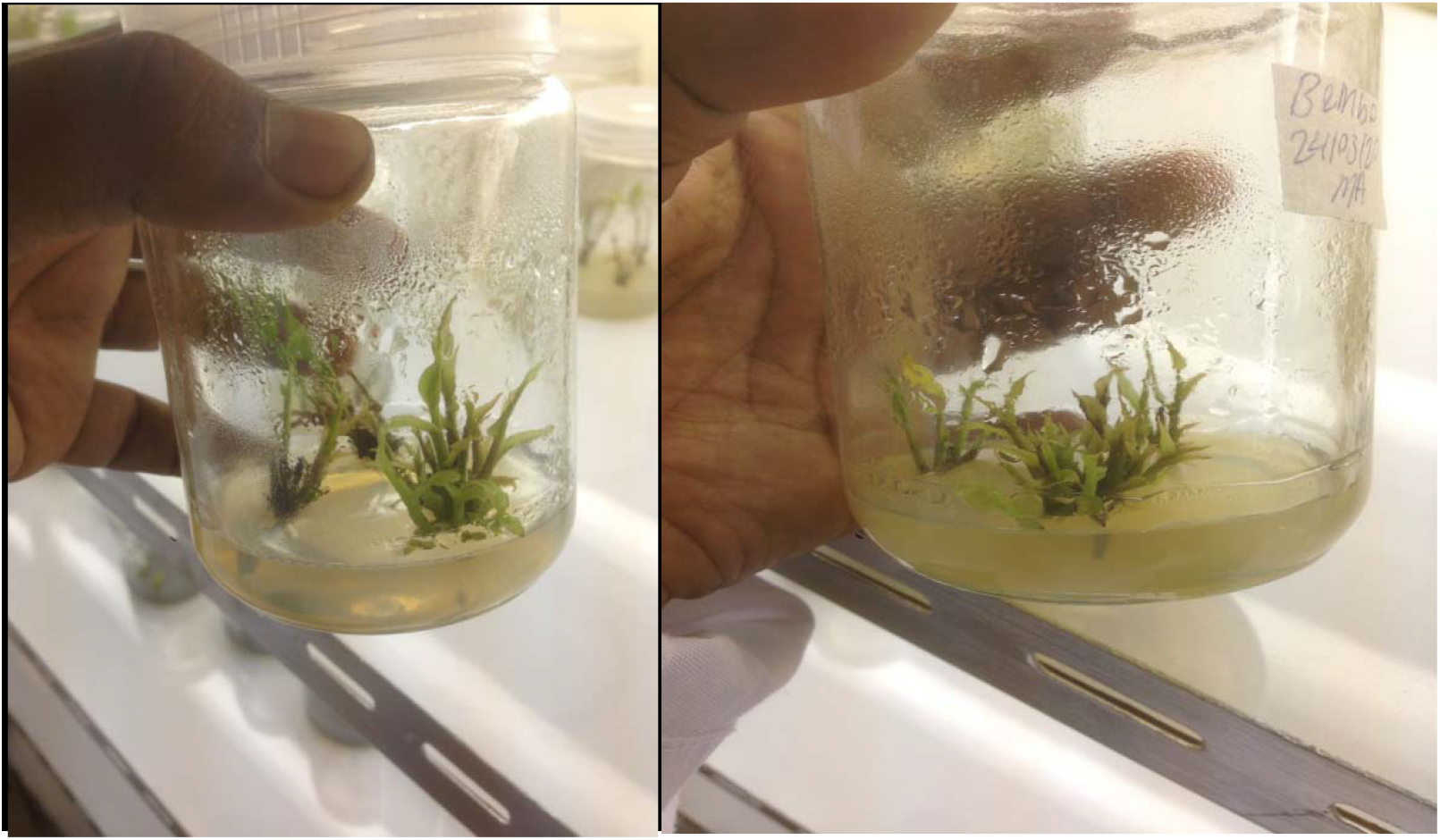
Multiplication of new leaf from intiated seed at 0.004 g /L BAP hormone.

### 2.7. Rooting of Shoots

The in vitro regenerated shoots, three shoots in abunch, were used for rooting studies after sub-cultured on PGRs free MS medium for 2 weeks. The rooting response of these shoots was studied on different concentrations of IBA (0.004, 0.005 and 0.006 g/L) and with for each treatmants used 0.1 g/L activated charkol for inhibition of oxidants of the cells mostlly for phenol exudation on MS medium and without hormone was used as control. For each treatment three jars, each with three clumps were used. All shoots were incubated on rooting medium for 4 weeks and kept under light conditions in culture room in CRD arrangement. Among all treatment, 0.005 g/L of IBA solution experment were best for root formation as it showen in **figure 7**.

**Figure 6.**
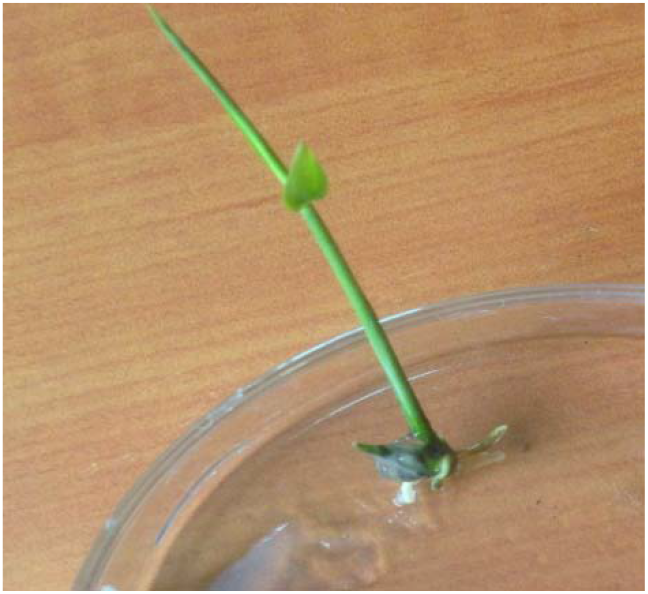
Root formation of Low land bamboo via IBA hormone after a week from single shoot.

**Figure 7.**
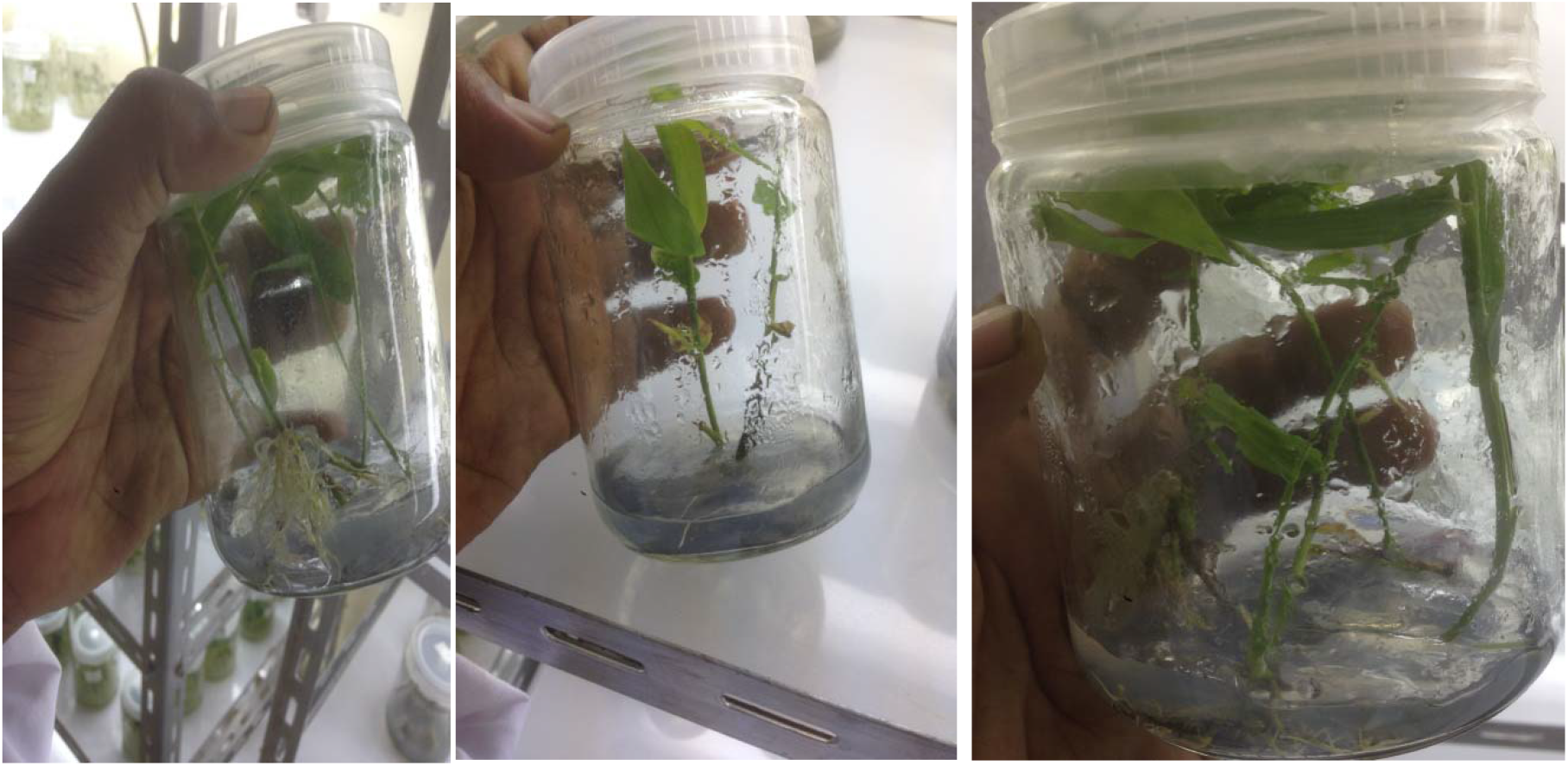
Root formation of Low land bamboo via IBA hormone after one month.

## 3. Results and discussion

### 3.1 Disinfection of the explants

For an effective micropropagation protocol and other applications of in −vitro cultures, explants must be disinfected at suitable disinfectant concentration for specified period where the explants can stay viable and contaminants free [25,26]. Therefore, in this experiment disinfection of bamboo seed were under took by 2 % (w/v) NaOCl solution for 25 min, 2-3 drops of Tween −20 for 20 minutes and antifungal of 20 g/l for 20 minute were the most effective disinfection treatment, which gave highest germination percentage; lowest contamination; and moderately clean explants.

### 3.2 Effect of BAP on establish-ment of culture shoots

In this experment all viable seeds were proliferated shoots after 5-7 days of culturing in both control and cytokinin fortified MS medium. How ever, the initiation percentage, the days for initiation, number of shoots initiated, length of shoots and leaves number were found vary in the different concentrations of cytokinins and control treatment. The best shoot initiation was recorded from 0.005 g/L 3-BAP supplemented in a MS medium. This showed that the shoot intiation percentage from seed was greatly influenced by types and concentrations of cytokinin. Their ability in this study for enhancing seed germination [27] and shoot initiation [28] is lined in this authors, investigations have been revealed that cytokinins were a key factor for bamboo species seed germination and multiple shoot proliferation [29]. Culturing of seeds for more than 30 days in a medium resulted browning of shoots and consequently died up the whole plantlet. Generally the present study indicated that the effect of 0.005 g /L BAP was best for shoot proliferation percentage and multiple shoot induction. The is research result is in agreement with the findings of other workers who have noted the effectiveness of BAP for the induction of multiple shoot from seeds in different bamboo species [30,31]. The longest (13.9 cm) and shortest (3.3 cm) shoots were recorded from 0.005 g/L −3 BAP and PGRs free fortified MS medium respectivel. Seeds cultured at MS medium supplemented from the present treatment higher BAP concentration induce greater number of shoots but without roots. This is due to the inapprporiate balance between cytokinins to auxin ratio in which the high level of cytokinins favors only shoot regeneration in the absence of equivalent auxin levels inside the bamboo plant.

### 3.3 Effect of BAP on shoot multiplication

Cytokinins were known to promote the function of other growth regulators like 2-isopentenyladenosine and zeatin [32]. In this study too, the addition of BAP on most microshoots of O. abyssinica resulted in an increased multiplication rate and higher mean shoot number over PGRs free MS medium. Investigated BAP at 0.005 g/L showed a good multiplication rate. The effect of BAP in inducing multiple shoots has already been reported in bamboo species like Arundinaria callosa [33] and Bambusa oldhamii [34]. Interestingly, the synergistic effect of BAP and KN for increased shoot multiplication rate and proliferation was also reported on Bambusa tulda and Melocanna baccifera [35]. The occurrence of phenol exudation at the cut ends of explants was the main problem faced during the multiplication study as showen in **Figure 8**. This phenolic exudation delayed the time required for sub-culturing accompanied with gradual browning of the shoots leaf and medium that eventually ends up in death. The IBA was found nice in both rooting percent and number of roots produced, which is in agreement with reports of Parthiban et al. and Diab and Mohamed [36,37].

**Figure 8.**
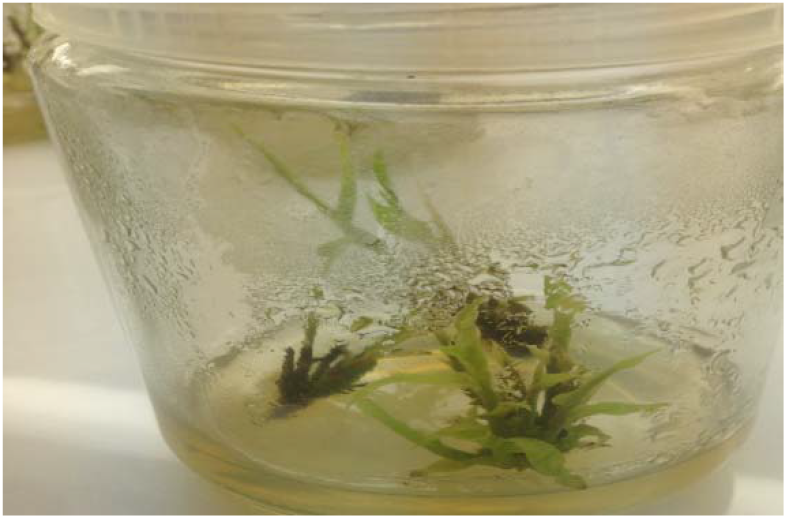
Browing of bamboo leaf due to phenol release.

## 4. Conclusions and recommendation

2 % of NaOCl solution for 25 min, 2-3 drops of Tween −20 for 20 minutes and antifungal of mancozine of 20 g/l for 20 minute were effective for disinfection of low land bamboo seed. 0.004 g/L −3 BAP supplementedwith MS medium showed best shoot proliferation, better shoot number and requires 5 −7 days to induce shoot. Similarly, for the shoot multiplication experiment, the tested cytokinin at 0.004 g/L BAP gave the efficient shoot number and shoot multiplication. In the root induction, IBA was also best at 0.005 g/L supplemented with MS medium gave best root number.

Finally,this study recommends to use this protocol for mass propagation of low land bamboo for reafforstation of degraded land which is highly exsposed for drought and for industery purpose.

## 5. Acknowledgements

I am highly acknowlede Ethiopian Enviroment, Forest and Climate Change Commisions for financing this research via UNDP Institutional Strengthening for the Forest Sector Development Program.

## References

1. BPG (Bamboo Phylogeny Group). An updated tribal and subtribal classification of the bamboos (Poaceae: Bambusoideae). In: Proceedings of the 9th World Bamboo Congress. Antwerp, Belgium: World Bamboo Organization. 2012;3–27.

2. Singh SR, Singh R, Kalia S, Dalal S, Dhawan AK, Kalia RK. Limitations, progress and prospects of application of biotechnological tools in improvement of bamboo-a plant with extraordinary qualities. Physiology and Molecular Biology of Plants. 2013;9(1):21–41.

3. Clayton, W.D., Vorontsova, M.S., Harman, K.T. and Williamson, H. (2006). Grass Base - The Online World Grass Flora. http://www.kew.org/data/grasses-db.html. [accessed 08 November 2015); 15:30 GMT.

4. Lobovikov M. (2005). Introduction to the Global Bamboo Resources Statistics, International Bamboo Inventory Training Workshop, 24 October –04 November 2005 in Beijing and Zhejiang Province, China

5. Ohrnberger, D. (1999). The bamboos of the world. Elsevier. 585 pp. Phillips, S. (1995). Flora of Ethiopia and Eritrea, Poacea (Gramineae) (Vol. 7). The National Herbarium, Addis Ababa, Ethiopia

6. Anonymous, (2016). https://en.wikipedia.org/wiki/List_of_bamboo-species.

7. Dwivedi A.P. (1993). Forests: The non-wood resources. International Book distributors, Dehra Dun, India. 352 pp.

8. Scurlock J. M.O., Dayton D.C. and Hames, B. (2000). Bamboo: an overlooked biomass resource. Biomass and Bioenergy 19: 229–244.

9. Wong K. M. (2004). Bamboo, the amazing grass: A guide to the diversity and study of bamboos in Southeast Asia. Selangor DarulEhsan, Malaysia: International Plant Genetic Resources Institute (IPGRI); and Kuala Lumpur, Malaysia: University of Malaya.

10. FAO (Food and Agriculture Organization). Global forest resources assessment. Main report. FAO Forestry Paper 163. Food and Agriculture Organization of the United Nations, Rome, Italy; 2010.

11. Seyoum K. Anatomical characteristics of Ethiopian low land bamboo (Oxytenanthera abyssinia). Presentation of the International Network for Bamboo and Rattan (INBAR). Beijing, China; 2008.

12. Yigardu, M., Asabeneh, A., Zebene, T., (2016). BAMBOO SPECIES INTRODUCED IN ETHIOPIABIOLOGICAL, ECOLOGICALAND MANAGEMENT ASPECTS, Ethiopian Environment and Forest Research Institute (EEFRI). 2016

13. Yiping L, Yanxia L, Henley KB, Guomo Z. Bamboo and climate change mitigation: a comparative analysis of carbon sequestration. International Network for Bamboo and Rattan (INBR), Technical Report. 2010;32.

14. Hall JB, Inada T. Oxytenanthera abyssinica (A.Rich. Munro). In: Louppe, D., Oteng-Amoako, A.A. & Brink, M. (Editors). Prota 7(1): Timbers/Bois d’œuvre 1. [CD-Rom] PROTA, Wageningen, Netherland; 2008.

15. Bartholomew IO, Maxwell EI. Phytochemical constituents and in vitro antioxidant capacity of methanolic leaf extract of Oxytenanthera abyssinica (A.Rich Murno) determined at varying concentrations of the plant extract. European Journal of Medicinal Plants. 2013;3(2):206–217.

16. Sisay F.. Site factor on nutritional content of Arundinaria alpine and Oxytenanthera abyssinica bamboo shoots in Ethiopia. Journal of Horticulture and Forestry. 2013;5(9):115–121.

17. Banik RL. Selection criteria and population enhancement of priority bamboos. INBAR Technical Report N°5 Genetic enhancement of bamboo and rattan INBAR New Delhi. 1995;99–110.

18. Bereket H. Study on establishment of bamboo processing plants in Amhara Regional State. M.Sc. Thesis Addis Ababa University, Addis Ababa, Ethiopia; 2008.

19. Demelash A, Zebene T, Yared K. Effect of storage media and storage time on germination and field emergence of Oxytenanthera abyssinica seeds. International Journal of Basic and Applied Sciences. 2012;1(3):218–226.

20. Kassahun E, Weih M, Ledin S, Christersson L. Biomass and nutrient distribution in a highland bamboo forest in southwest Ethiopia: Implications for management. Forest Ecology and Management. 2005;204:159–169.

21. Alexander MP, Rao TC. In vitro culture of bamboo embryo. Current Science. 1968;37:415.

22. Arya S, Sharma S, Kaur R, Arya ID. Micropropagation of Dendrocalamus asper by shoot proliferation using seeds. Plant Cell Report. 1999;18:879–882.

23. Arya ID, Kaur B, Arya S. Rapid and mass propagation of economically important bamboo Dendrocalamus hamiltonii. Indian Journal of Energy. 2012;1(1):11–16.

24. Devi WS, Bengyella L, Sharma GJ. In vitro seed germination and micropropagation of edible bamboo Dendrocalamus giganteus Munro using seeds. Biotechnology. 2012;1–7.

25. Murashige T, Skoog F. A revised medium for rapid growth and bioassays with tobacco tissue cultures. Physiol Plant. 1962;15:473–97.

26. Kulus D. Micropropagation of Kalanchoe tubiflora (Harvey) Hamet. Nauka, Przyroda, Technologie. 2015;9(1):1–8.

27. Wegayehu F, Firew M, Belayneh A. Optimization of explants surface sterilization condition for field grown peach (Prunus persica L. Batsch. Cv. Garnem) intended for in vitro culture. African Journal of Biotechnology. 2015;14(8):657–660.

28. Oyebanji OB, Nweke O, Odebunmin NB, Galadima NB, Idris MS, Nnod UN, Afolabi AS, Ogbadu GH. Simple, effective and economical explant surface sterilization protocol for cowpea, rice and sorghum seeds. African Journal of Biotechnology. 2009;8(20):5395–5399.

29. Teixeira da Silva, Jaime A, Kulus D, Zhang X, Zeng S, Ma G, Piqueras A. Disinfection of explants for saffron (Crocus sativus L.) tissue culture. Environmental and Experimental Biology. 2016;14(4):183–198.

30. Miransaria M, Smith DL. Plant hormones and seed germination: Review. Environ-mental and Experimental Botany. 2014;99:110–121.

31. Ashraf MF, Aziz M, Kemat N, Ismail I. Effect of cytokinin types, concentrations and their interactions on in vitro shoot regeneration of Chlorophytum borivilianum Sant. & Fernandez. Electronic Journal of Biotechnology. 2014; EJBT–00047:1–5. Available: http://dx.doi.org/10.1016/j.ejbt.2014.08.004 Accessed on September 16, 2014

32. Nadgir AL, Phadke CH, Gupta PK, Parsharami VA, Nair S, Mascarenhas AF. Rapid multiplication of bamboo by tissue culture. Silvae Genetica. 1984;33(6):219–223.

33. Tuan TT, Tu HL, Giap DD, Du TX. The increase in in vitro shoot multiplication rate of Dendrocalamus asper (Schult. f.) Back. ex Heyne. TAP CHÍ SINH HOC. 2012;34(3se):257–264.

34. Woeste KE, Vogel JP, Kieber JJ. Factors regulating ethylene biosynthesis in etiolated Arabidopsis thaliana seedlings. Physiologia Plantarum. 1999;105:478–484.

35. Saikia SP, Mudoi KD, Borthakur M. Effect of nodal positions, seasonal variations, and shoot clump and growth regulators on micropropagation of commercially important bamboo, Bambusa nutans Wall. Ex. Munro. African Journal of Biotechno-logy. 2014;13(9):1961–1972.

36. Gaspar T, Kevers C, Penel C, Greppin H, Reid DM, Thorpe TA. Plant hormones and plant growth regulators in plant tissue culture: Review. In vitro Cellular and Developmental Biology Plant. 1996;32: 272–28.

37. Zaerr JB, Mapes MO. Action of growth regulators. In: Bonga, J.M. Durzan, D.J. (Eds.). Tissue Culture in Forestry. Martinus Nijhoff/Dr. W. Junk Publisher, The Hague/Boston/London. 1982;231–255.

38. Devi WS, Sharma GJ. In vitro propagation of Arundinaria callosa Munroan edible bamboo from nodal explants of mature plants. The Open Plant Science Journal. 2009;3:35–39.

